# Study of Real-Valued Distance Prediction For Protein Structure Prediction with Deep Learning

**DOI:** 10.1101/2020.11.26.400523

**Authors:** Jin Li, Jinbo Xu

**Affiliations:** Toyota Technological Institute at Chicago; Department of Computer Science, University of Chicago

## Abstract

Inter-residue distance prediction by deep ResNet (convolutional residual neural network) has greatly advanced protein structure prediction. Currently the most successful structure prediction methods predict distance by discretizing it into dozens of bins. Here we study how well real-valued distance can be predicted and how useful it is for 3D structure modeling by comparing it with discrete-valued prediction based upon the same deep ResNet. Different from the recent methods that predict only a single real value for the distance of an atom pair, we predict both the mean and standard deviation of a distance and then employ a novel method to fold a protein by the predicted mean and deviation. Our findings include: 1) tested on the CASP13 FM (free-modeling) targets, our real-valued distance prediction obtains 81% precision on top L/5 long-range contact prediction, much better than the best CASP13 results (70%); 2) our real-valued prediction can predict correct folds for the same number of CASP13 FM targets as the best CASP13 group, despite generating only 20 decoys for each target; 3) our method greatly outperforms a very new real-valued prediction method DeepDist in both contact prediction and 3D structure modeling; and 4) when the same deep ResNet is used, our real-valued distance prediction has 1-6% higher contact and distance accuracy than our own discrete-valued prediction, but less accurate 3D structure models.

## Introduction

Immense progress has been made on protein structure prediction due to the application of deep ResNet (convolutional residual neural network) that can accurately predict inter-residue or inter-atom relationships[1]–[6]. These predicted outputs, such as inter-residue contact or distance, are key to currently the most successful structure prediction methods[7]. Early contact prediction algorithms like MetaPSICOV use traditional machine learning methods to predict contacts individually[8]. However, this is suboptimal because they predict contacts between two atoms regardless of the other atoms. To address this, RaptorX introduced a deep ResNet to predict all contacts of a protein (or a big chunk) simultaneously[2]. Deep ResNet is able to learn complex sequence-structure relationships and make use of high-order contact correlation to achieve much better accuracy. Right after its success on contact prediction, RaptorX moved to distance prediction by ResNet because distance conveys more information for structure modeling[5], [9]. Distance prediction is also adopted by AlphaFold[3], a leading method in CASP13. However, both RaptorX and AlphaFold discretize distance into dozens of bins and formulates the distance prediction problem as a multi-class classification problem.

Alternatively, as suggested in the RaptorX paper[5], it is also possible to predict real-valued distance by deep learning. A natural question to ask is how well real-valued distance can be predicted and how useful it is for 3D structure modeling. DeepDist[10] is one of the very few methods that predict real-valued distance by ResNet. DeepDist is very complex, as it trains multiple deep models of different architectures on different types of data. For 3D structure modeling, DeepDist converts real-valued prediction to distance bounds and then feeds them into CNS[11], a software designed for experimental structure determination. However, DeepDist did not report how well real-valued distance prediction alone can fold a protein, but only showed that protein folding may be improved by adding real-valued prediction on top of discrete-valued prediction. Ding et al. developed another real-valued prediction method by adding generative adversarial networks (GANs) on top of ResNet to enforce global distance consistency[12]. Similar to DeepDist, Ding et al. also derived distance bounds from real-valued prediction, which are then fed into CNS for 3D structure modeling. However, even with recent progress in improving the stability of GANs, they are still notoriously difficult to train and scale to larger networks[13]. Neither DeepDist nor Ding et al’s work have compared their real-valued prediction to discrete-valued prediction based upon the same deep network and thus, cannot accurately evaluate the strength and weakness of real-valued distance prediction compared to discrete-valued prediction.

In this paper, we present a new method for real-valued prediction of inter-atom distance and inter-residue orientation. Our method differs from DeepDist and Ding et al’s method in that we predict both mean and standard deviation (i.e., a normal distribution) of distance and orientation while they predict only a single value (which can be interpreted as mean). Our prediction pipeline is much simpler than DeepDist and easier to train than GANs. We also introduce a novel way of using the predicted mean and deviation to build 3D models that is distinct from how discrete distance is used. Our experimental results show that our real-valued prediction exceeds the best in CASP13 in terms of both contact accuracy and 3D structure modeling and that our method greatly outperforms DeepDist and compares favorably to Ding et al’s method. Finally, we will rigorously compare real-valued prediction to discrete-valued prediction based upon the same deep network, which is missing in both DeepDist and Ding et al’s work. We find that when the same deep ResNet is used, real-valued prediction has higher contact and distance accuracy, but less accurate 3D models than discrete-valued prediction.

## Results

### Overview of the method

Our real-valued prediction method consists of two steps 1) predicting the mean and standard deviation of backbone conformation attributes by a deep ResNet. We simultaneously predict the distance of backbone atom pairs (Cb-Cb, Ca-Ca, and N-O) and inter-residue orientation angles defined in trRosetta[4]; 2) fitting the harmonic function with the predicted mean and deviation as constraints to build 3D structure models by gradient descent. We use PyRosetta to build 3D models by performing gradient-based minimization and then the fast relaxation protocol to pack side chains and reduce steric clashes[14]. In contrast, our discrete-valued prediction uses a spline function to construct distance and orientation potential for gradient-based minimization[15].

For both real-valued and discrete-valued predictions, we train six deep ResNet models of the same architecture on the same training data and ensemble them to make predictions. Please see [5] for a detailed description of our ResNet network, but here we use a much larger ResNet. In particular, our current ResNet has ~60 ResNet blocks, each consisting of two 2D convolution layers and 2 batch normalization layers. On average, each convolutional layer has ~170 filters and in total a deep ResNet model has ~50 million parameters. We use mixed-precision training to reduce the training time and GPU memory usage by half without losing accuracy[16]. The ResNets for our real-valued and discrete-valued predictions are the same except the output layer. For real-valued prediction, the output layer generates two values for an atom (or residue) pair, representing predicted mean and standard deviation. For discrete-valued prediction, the output layer yields one value for each discrete bin, representing its predicted probability.

We use the following input features for our deep ResNet: 1) primary sequence represented by one-hot encoding; 2) sequence profile derived from multiple sequence alignments (MSA) that encode evolutionary information at individual residues. We also use secondary structure and solvent accessibility predicted from sequence profile; 3) co-evolution information including mutual information and the output matrices generated by CCMpred[17].

### Accuracy of predicted contacts on CASP13 FM and FM/TBM targets

Using the predicted mean and deviation, we may estimate the probability of two residues forming a contact (i.e., having distance<8Å). As shown in Table 1, our real-valued contact prediction has slightly better accuracy than our discrete-valued prediction and both of them outperform the best methods in CASP13 by a good margin. Our real-valued method greatly outperforms DeepDist, a very new method for real-valued distance prediction. While our top L/5, L/2, and L contact precisions (L is sequence length) for the 43 FM and FM/TBM targets are 84.6%, 72.6%, and 61.8%, DeepDist’s contact precisions are 78.6%, 64.5%, and 49.6% [10]. Our methods also have better contact precision than trRosetta[4], a method developed after CASP13 that employs discrete-valued prediction, although trRosetta used a newer sequence database uniclust30 (dated in August 2018) and a larger metagenome database to generate MSAs (multiple sequence alignment) than us.

**Table 1.** Precision and F1 (%) of long-range contact prediction of FM and FM/TBM CASP13 targets by several competing methods. The F1 of AlphaFold is taken from [5]. The trRosetta result is taken from [4].

|  | 31 CASP13 FM targets |  |  | 12 CASP13 FM/TBM targets |  |  |
| --- | --- | --- | --- | --- | --- | --- |
|  | Top L/5 | Top L/2 | Top L | Top L/5 | Top L/2 | Top L |
|  | F1 of long-range contact prediction |  |  |  |  |  |
| AlphaFold in CASP13 | 22.7 | 36.9 | 41.9 | 31.4 | 48.7 | 55.1 |
| RaptorX in CASP13 | 23.3 | 36.2 | 41.1 | 28.8 | 43.2 | 51.8 |
| Zhang in CASP13 | 21.2 | 34.1 | 39.2 | 28.4 | 43.3 | 49.5 |
| Discrete (our work) | 27.5 | 44.1 | 51.1 | 31.9 | 52.9 | 62.0 |
| Real-Valued (our work) | 27.9 | 44.6 | 51.8 | 31.8 | 52.4 | 62.4 |

|  | Precision of long-range contact prediction |  |  |  |  |  |
| --- | --- | --- | --- | --- | --- | --- |
| RaptorX in CASP13 | 70.0 | 58.0 | 45.0 | 85.8 | 70.1 | 56.9 |
| Zhang in CASP13 | 65.7 | 54.8 | 39.1 | 82.3 | 70.0 | 54.8 |
| trRosetta | 78.5 | 66.9 | 51.9 | NA | NA | NA |
| Discrete (our work) | 80.2 | 67.6 | 57.6 | 93.9 | 83.5 | 70.7 |
| Real-Valued (our work) | 81.2 | 68.7 | 58.1 | 93.6 | 82.8 | 71.3 |

#### Real-valued vs. discrete-valued prediction

We use the same hyperparameters that are optimized on the discrete-valued ResNet for our real-valued ResNet. Our real-valued prediction still produces marginally better (0.5-1.0%) top L long-range contact accuracy than our discrete-valued prediction. When the extra long-range contact prediction is evaluated, our real-valued method achieves a top L/5, L/2, and L precision of 65.5%, 55.4%, and 49.2%, respectively, whereas our discrete-valued prediction has precision 63.1%, 53.6% and 47.8%, respectively. We say one contact is extra long-range if its two involving residues are separated by at least 48 residues along the primary sequence. If we were to more vigorously tune the hyperparameters for our real-valued deep ResNet models, our results would be even better. There is a high correlation (CC=0.98) between our discrete-valued and real-valued L/5 contact precision on the 43 hard targets, but there are still 9 test proteins with precision difference greater than 5%, indicating that there are situations where real-valued prediction may be more useful (see Figure 1).

**Figure 1.**
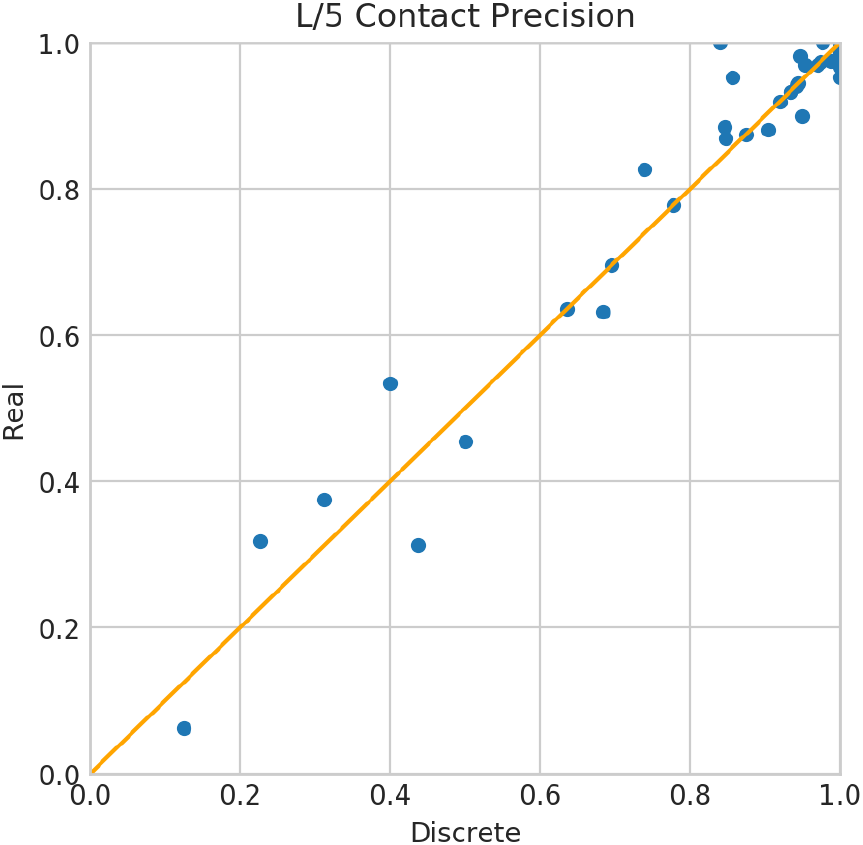
Top L/5 contact precision of our discrete-valued and real-valued methods on the 43 FM and FM/TBM CASP13 targets. A dot above the diagonal line indicates that real-valued prediction is better than its corresponding discrete-valued prediction.

### Distance prediction accuracy on CASP13 FM and FM/TBM targets

We use a few metrics to evaluate distance accuracy of our real-valued and discrete-valued predictions, including absolute error, relative error, precision, recall, F1, pairwise distance test (PDT), and high-accuracy pairwise distance test (PHA), which are explained in section Methods. While evaluating distance prediction, we consider only those long-range atom pairs with predicted distance <15Å and native distances <15Å. To evaluate our discrete-valued prediction, we convert discrete probability distributions to real-valued distance by computing the expected value of a discrete distribution, which is detailed in [5]. As shown in Table 2 and Appendix Fig. S1, our real-valued distance prediction is better than our own discrete-valued prediction by 1-6% in terms of all the metrics except recall. We do not compare our distance accuracy with Ding et. al.’s GAN method because they only reported distance accuracy on their validation set but not on the CASP13 targets. In addition, because it is not clear how DeepDist precisely defines their distance measures, it is challenging for us to compare our distance accuracy with DeepDist, but we have much better contact accuracy than DeepDist.

**Table 2.** Average distance accuracy of our real-valued and discrete-valued prediction methods. The accuracy is measured by absolute error, relative error, precision, recall, F1, PHA and PDT.

|  | CbCb |  | CaCa |  | NO |  |
| --- | --- | --- | --- | --- | --- | --- |
|  | Real | Discrete | Real | Discrete | Real | Discrete |
| Abs. Error | 4.069335 | 4.232118 | 3.759566 | 3.974853 | 3.643687 | 3.872508 |
| Rel. Error | 0.241389 | 0.251571 | 0.220723 | 0.232381 | 0.215353 | 0.227033 |
| Precision | 0.682059 | 0.650375 | 0.691564 | 0.670452 | 0.692480 | 0.676140 |
| Recall | 0.712129 | 0.736852 | 0.728711 | 0.745034 | 0.748851 | 0.759924 |
| F1 | 0.686266 | 0.679169 | 0.698683 | 0.694299 | 0.710045 | 0.705230 |
| PHA | 0.431603 | 0.412376 | 0.462826 | 0.444350 | 0.481869 | 0.464568 |
| PDT | 0.610425 | 0.589494 | 0.637133 | 0.617010 | 0.650349 | 0.630967 |

### Accuracy of predicted 3D models on CASP13 FM and FM/TBM targets

We use the predicted real-value attributes to generate only 20 decoys for each target and then select 5 decoys with the lowest energy as our prediction of 3D models. On the 32 CASP13 FM targets, the average quality (measured by TMscore[18]) of the first and best (of 5) models is 0.582 and 0.599, respectively. On the 13 FM/TBM targets, the average TMscore of the first and best (of 5) models is 0.641 and 0.651, respectively. When the best models are considered and TMscore>0.5 is used to judge if a predicted 3D model has a correct fold or not, our real-valued method predicts correct folds for 23 of the 32 FM targets and 11 of the 13 FM/TBM targets. Despite generating only 20 decoys per target, our real-valued method performs as well as AlphaFold in CASP13, which has an average TMscore 0.583 for the first models of the 32 FM targets and correctly folds 23 of the 32 targets[3]. It is reported that AlphaFold generated thousands of decoys per target.

DeepDist reported an average TMscore 0.487 and 0.522 for the first and best models of the 43 CASP13 hard (FM and FM/TBM) targets, respectively. In total DeepDist predicted correct folds for 23 of the 43 targets (see Table 3). In contrast, our real-valued prediction method obtains an average TMscore of 0.604 and 0.619 for the first and best models, respectively, and predicts correct folds for 33 of the 43 targets. Our method also vastly outperforms another distance-based folding method DMPfold[19], which has average TMscore 0.438 for the first models. Ding et. al. evaluated their real-valued prediction on only 20 FM and FM/TBM targets and reported an average TMscore of 0.620, whereas we can achieve a similar TMscore of 0.612 with real-valued prediction[12], [19]. However, the comparison with Ding et al.’s result is not rigorous, as they used the official domain sequences as inputs while we do not. To simulate the real-world prediction scenarios, we predicted 3D models on the domains determined by our server during the CASP13 season in the absence of the native structures of the test proteins. When evaluating the quality of our predicted 3D models, we only count the segments that overlap with the official domains. As such, when our own domain definition deviates significantly from the official one, our predicted 3D models have a low quality score even if we may build a good 3D model for our own domain.

**Table 3.** Average TMscore obtained by 3 competing methods on the 43 CASP13 FM and FM/TBM targets. The DeepDist and DMPfold results are taken from the DeepDist paper[10].

| Method | Top 1 | Top 5 | #Correct Folds |
| --- | --- | --- | --- |
| Ours | 0.604 | 0.619 | 33 |
| DeepDist | 0.487 | 0.522 | 23 |
| DMPfold | 0.438 | 0.449 | 16 |

#### Real-valued vs. discrete-valued prediction

Similarly, we also generate 20 decoys for each target using our discrete-valued prediction and select the top 5 decoys for each target by energy. On the 32 CASP13 FM targets, the average TMscore of the first and best models generated by our discrete-valued prediction is 0.646 and 0.672, respectively. On the 13 FM/TBM targets, the average TMscore of the first and best 3D models is 0.671 and 0.683, respectively. Our discrete-valued method can predict correct folds for 26 of the 32 FM targets and 11 of the FM/TBM targets. That is, although our real-valued prediction generates better contact accuracy, its 3D modeling accuracy is not as good as our discrete-valued prediction. The correlation between our real-valued 3D model quality (measured by TMscore) and discrete-valued model quality is 0.95, but our discrete-valued method predicts better 3D models for nearly all targets (Figure 2). The correlation between the top L/2 contact precision of the 31 FM targets with their first model TMscore is 0.626 for discrete-valued prediction and 0.543 for real-valued prediction. That is, there is still room for improvement in real-valued-based 3D structure modeling. This also implies that contact precision is a better indicator for 3D model quality of discrete-valued prediction than real-valued prediction. The correlation between the logarithm of MSA depth (i.e., ln(Meff)) and model quality for real-valued and discrete-valued predictions is 0.572 and 0.557, respectively. When ln(Meff)>4.0, our discrete-valued method can predict the correct folds for all targets while our real-valued method fails on one target (Appendix Fig. S2).

**Figure 2.**
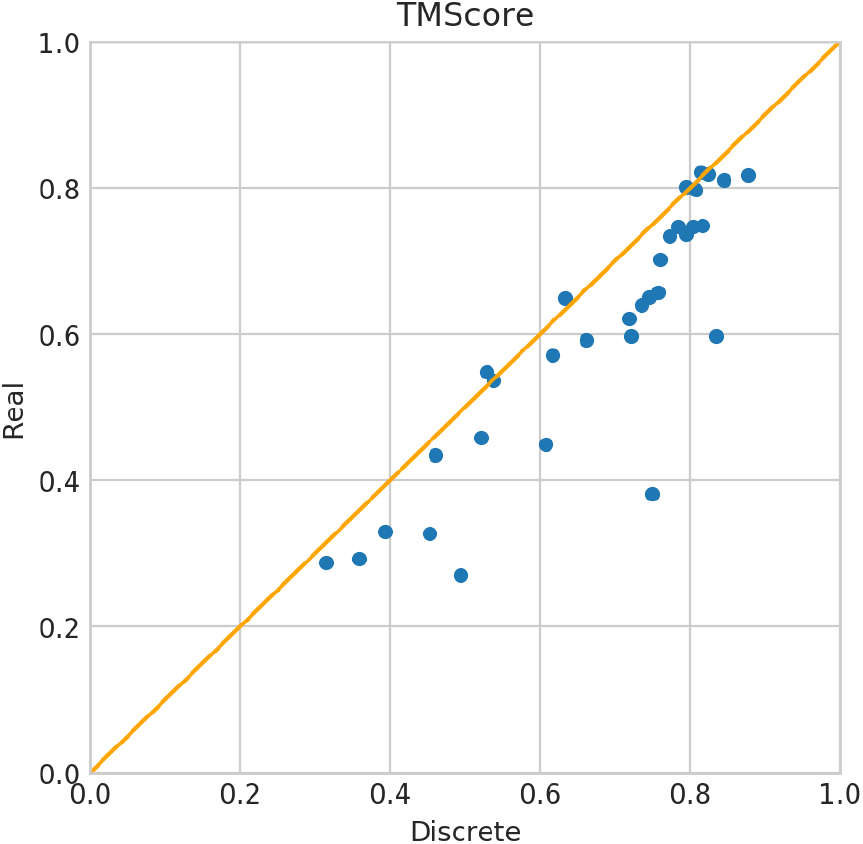
TMScore of the first models of our discrete-valued vs real-valued predictions for the 32 CASP13 FM targets.

### Strength and weakness of real-valued prediction

Real-valued prediction has both advantages and down-sides. First, dozens of parameters are needed to represent a discrete distance distribution while only two parameters (mean and standard deviation) are used for real-valued prediction. Second, discretizing distance possibly reduces the amount of information that can be learned by a machine learning method. Indeed, our experiments showed that real-valued prediction has slightly better contact and distance accuracy. Because we model the energy function for real-valued prediction using the harmonic function that has only two parameters, our real-valued energy function is symmetric across the mean, much smoother, and much simpler than our discrete-valued potential. As shown in Appendix Fig. S3, discrete distance potentials have troughs followed by peaks followed by another through, which is an undesirable characteristic for energy minimization. The smoothness of real-valued energy function makes gradient-based minimization easier. However, discrete-valued prediction uses dozens of parameters to define a probability distribution and thus, result in a higher resolution energy function for 3D structure modeling than real-valued prediction. This shall be the major reason why our real-valued prediction underperforms in 3D structure modeling.

### Case study

Our real-valued and discrete-valued methods perform differently on two CASP13 hard targets T0990-D1 and T1008-D1. For T0990-D1, our discrete-valued prediction works better, while for T1008-D1 our real-valued prediction works better.

T0990-D1 has 76 residues and the logarithm of its MSA depth is 3.308. Our real-valued and discrete-valued predictions have similar contact precision. The top L/5, L/2, and L long-range contact precision of both methods are 0.933, 0.605, and 0.6. The top L/5 short-range contact precision is 0.733. But our real-valued prediction has better top L/5 medium-range contact precision (0.8) than discrete-valued prediction (0.667). Both methods have slightly different distance accuracy. Our discrete-valued prediction has a smaller relative and absolute distance prediction error (0.124, 1.424) than our real-valued prediction (0.145, 1.624). Our discrete-valued prediction also has slightly better PDT and PHA (0.794, 0.619) than our real-valued prediction (0.764, 0.580). However, our discrete-valued prediction produces much better 3D models than real-valued prediction in terms of TMscore (0.75 vs 0.382), RSMD (2.512 vs 9.819), GHA (0.536 vs 0.276), and GDT (0.75 vs 0.418). This suggests that contact accuracy may not necessarily be a good predictor of 3D model quality as it does not capture the overall information of the predicted distance map. In addition, the real-valued prediction may not necessarily predict better distance than the discrete-valued method.

**Figure 3.**
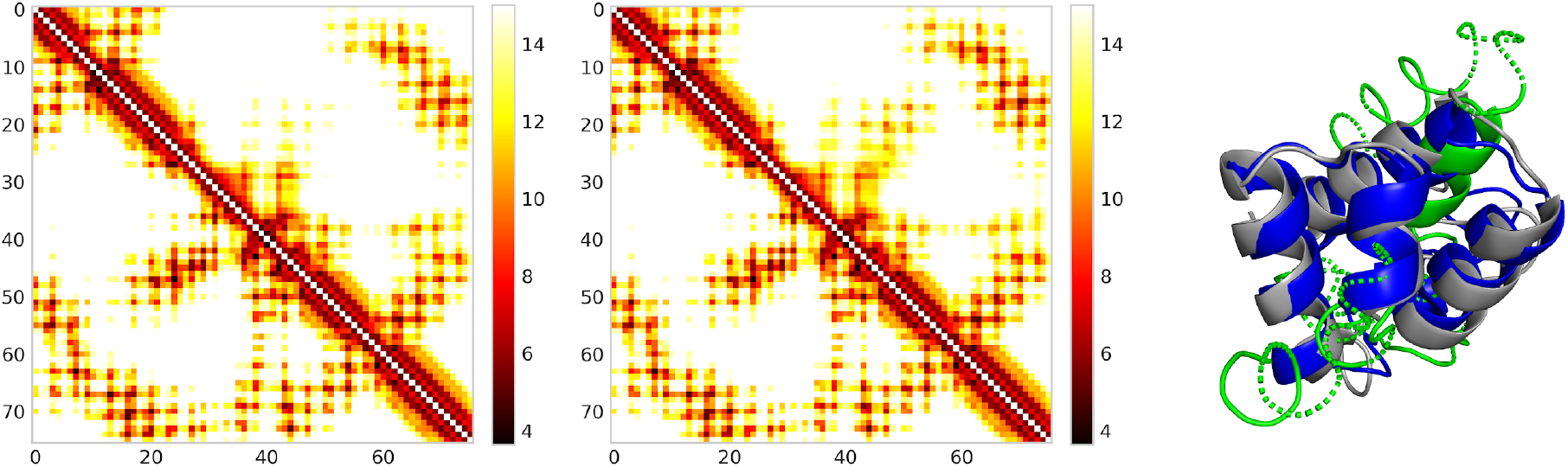
Distance map for T0990-D1 predicted by real-valued ResNet (Left) and discrete-valued ResNet (Middle). Only distances less than 15Å are displayed in colors. In each picture, native and predicted distance is shown below and above the diagonal line, respectively. The right picture shows the superimposition of T990-D1 native structure (gray), real-valued model (green), and discrete-valued model (blue).

For T1008-D1, our discrete-valued prediction has top L/5, L/2, and L long-range contact precision 0.933, 0.684, and 0.508, respectively, better than our real-valued prediction ( 0.933, 0.631, and 0.492). Our real-valued prediction has better predicted distance accuracy in terms of relative error (2.633 vs 3.014), absolute error (0.213 vs 0.231), PDT (0.608 vs 0.574), and PHA (0.385 vs 0.352). The 3D model built from our real-valued prediction is better across all metrics, including TMscore (0.587 vs 0.433), GDT (0.444 vs 0.416), RMSD (3.671 vs 7.923), and GHA (0.393 vs 0.312). This suggests that an improved distance prediction can help improve 3D structure modeling.

**Figure 4.**
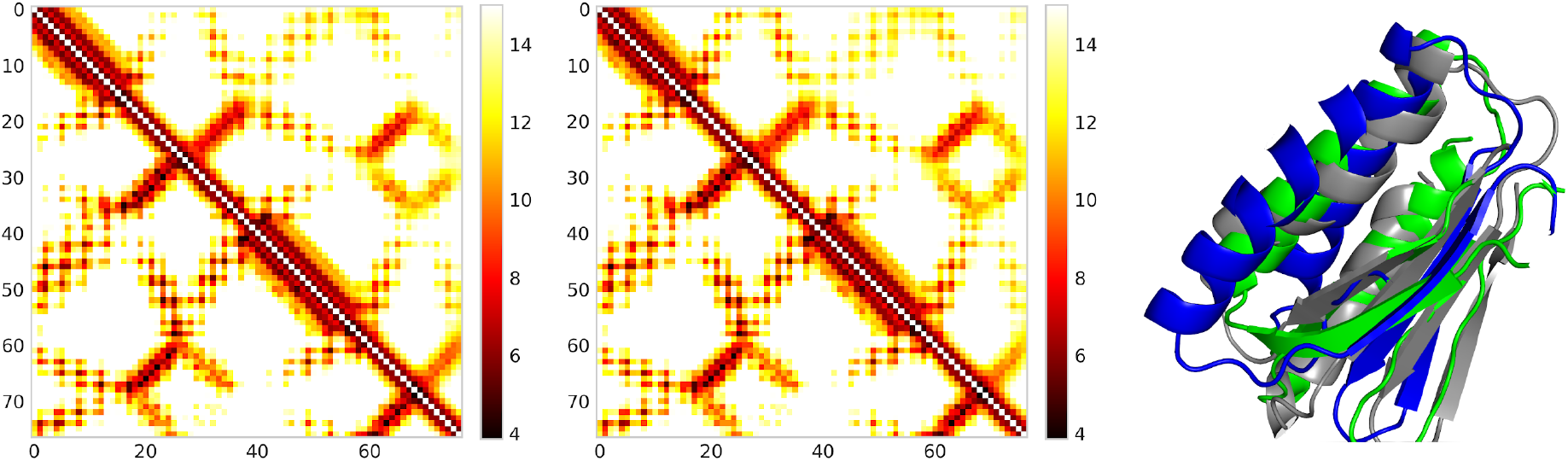
Distance map for T1008-D1 predicted by real-valued ResNet (Left) and discrete-valued ResNet (Middle). Only distances less than 15Å are displayed in colors. In each picture, native and predicted distance is shown below and above the diagonal line, respectively. The right picture shows the superimposition of T1008-D1 native structure (gray), real-valued model (green), and discrete-valued model (blue).

## Conclusion and Discussions

We have presented a new method for real-valued distance prediction using a deep ResNet. This method can achieve a top L/5 contact precision of 81.2%, more than 10% greater than the best methods in CASP13. Even generating only 20 decoys per target, our method can correctly fold the same number of CASP13 FM targets as the best human group in CASP13. Our method also outperforms existing real-valued prediction methods such as DeepDist in terms of both contact accuracy and 3D model quality. When the same ResNet is used, our real-valued prediction can achieve a 1-6% improvement over its discrete version in terms of contact and distance accuracy, but it falls short in 3D structure modeling accuracy. Even though the energy function predicted by our real-valued method is smoother and more symmetric, its resolution is not as high as our discrete-valued prediction.

There is still much room to improve real-valued prediction. For example, DeepDist trains both real and discrete predictions at the same time and then combines the results, which can further improve contact accuracy and folding ability. Ding et al. show that using the GAN on top of the deep ResNet can further improve the global consistency of distance prediction, which is something worth trying. It is also possible that we can employ better loss functions to deal with the class imbalances for protein distance prediction[20]. Recently, there has been much progress in making convolution layers more efficient and powerful, and it would be interesting to see how this can improve the ResNet architecture[21]–[25]. There has also been interest in end-to-end training for protein structure prediction, which can improve the learned relationship between structure and output while speeding up prediction time[26]–[28]. And lastly, we would continue to investigate ways in which we can reduce the gap between real-valued and discrete-valued prediction in 3D structure modeling.

## Methods

### Training and validation data

To conduct our experiments, we train and validate our models on the Cath S35 protein dataset downloaded in December 2019, which is a set of 32511 CATH domains (https://www.cathdb.info/) in which two domains share at most 35% sequence identity. We remove short domains (<25 residues) and those with too many missing Ca and Cb atoms. For each protein domain in Cath S35, we generate their multiple sequence alignments (MSAs) by running HHblits with E-value=0.001 on the uniclust30 library dated in October 2017 and then derive input features[29], [30]. We randomly split the dataset into a train and validation set (1800 domains). We generate 6 splits and train deep ResNets on each split. As shown in Xu et. al [31], there is very little difference between this Cath S35 dataset and the version dated in March 2018. The deep ResNet models trained on them have almost the same (contact prediction and 3D modeling) performance on the CASP13 FM and FM/TBM targets.

### Independent test data

We use the 45 CASP13 hard targets (32 FM targets and 13 FM/TBM targets) to evaluate all methods. Since T0953s1 and T0955 have very few long-range contacts, they are not used to evaluate contact or distance accuracy. We use HHblits (with E-value=0.01) and TMalign to check sequence profile and structural similarity between the CASP13 FM targets and our training set. When searching through our training set, HHblits returns a large E-value (>10) for most of the 32 FM targets. The only exceptions are targets T0975 and T1015s1; T0975 is related to 4ic1D (HHblits E-value=4.2E-12) and T1015s1 is related to 4iloA (HHblits E-value=0.024). However, since both 4ic1D and 4iloA were deposited to PDB well before 2018 and the structure similarity (as measured by TMscore) between them is both less than 0.5, it is fair to include them into our training set.

### MSA generation and input features

To generate multiple sequence alignments (MSAs) for the test targets, we run HHblits[29] with E-value=1E-3 and 1E-5 on the uniclust30 database dated in October 2017 and jackhammer[32] with E-value=1E-3 and 1E-5 on the uniref90 database dated in March 2018. If any target has a shallow MSA depth (ln(Meff)<6), we also search a metagenome database dated in June 2018 to see if more sequence homologs can be found. We chose these databases because they ensured fairness for comparisons, as they were created before the start of CASP13. The input features include both sequential and pairwise features. For sequential features, we use 1) the primary sequence represented as a one-hot encoding; 2) sequence profiles derived from MSAs that encoded evolutionary information at each residue; 3) secondary structure and solvent accessibility predicted from the sequence profile. Our pairwise features include coevolution information and consist of 1) mutual information; 2) CCMpred[17] output. The CCMpred output includes one L×L co-evolution matrix and one full precision matrix of dimension L×L×21×21 where L is the protein sequence length.

### Protein structure representation

We represent protein backbone conformation using inter-atom distance matrices (Ca-Ca, Cb-Cb, and N-O) and inter-residue orientation matrices as employed by trRosetta[4]. When training the real-valued ResNet models, all distances greater than 20Å are set to 20Å. If we do not do this, our model focuses on learning large distances instead of more useful small distances (<16Å). We tried setting the distance limit to 16Å instead of 20Å, but did not obtain better performance. We normalize distances and angles to be bounded by 0 and 1 and the dihedral angles to be bounded by −1 and 1. This normalization not only allows the gradients to flow more easily in the deep ResNet, but it also equally weights the loss function across the inter-atom predictions. For discrete-valued prediction, we discretize distance into 47 bins: 0-2Å, 2-2.4Å, 2.4-2.8Å,…,19.6-20Å, and >20Å.

### Deep model training

We follow AlphaFold and trRosetta’s technique of subsampling MSAs. During subsampling, we randomly sample 40-60% of the sequence homologs from an MSA (with at least 2 sequences) and then derive input features from the sampled MSA. To save training time, we generated 10 different samples on disk so that our training program just needs to randomly select one sample from the disk during training. Our deep ResNet learns to predict the mean and standard deviation of a normal distribution for real-valued distances and orientations. Because it is challenging for the deep ResNet to learn the mean and standard deviation simultaneously, we first train it to learn the mean and then the standard deviation while fixing the mean. We use the AdamW[33] optimizer with β1 set to 0.1, β2 set to 0.001, and L2 regularization factor of 0.35. We train our deep ResNet to predict the mean for 20 epochs with a learning rate of 0.0002, 1 more epoch with a learning rate of 0.00004, and the last epoch with a learning rate of 0.000008. We train the standard deviation parameters similarly, but for only 10 epochs because it converges much more quickly. We train up to 6 deep ResNet models with the same hyperparameters on different subsets of our data and ensemble them to predict the final mean and standard deviation.

Instead of predicting a normal distribution for the dihedral angles, we tried to predict the von Mises distribution[34]. However, because we train our models to predict distances and orientations simultaneously and it is hard for our models to learn with different loss functions at the same time, those models did not converge well. We also tried to train our models to predict discrete-valued and real-valued distance at the same time but failed for similar reasons.

### Building protein 3D models and model clustering

We build our 3D models from distance and orientation prediction with PyRosetta as follows: 1) convert the predicted mean μ and deviation σ to energy potential by fitting them to the harmonic function. That is, the potential of an orientation angle *x* is 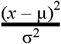 and the potential of a distance d function is 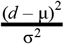 if d is <19, otherwise 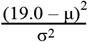. (2) minimize the energy potential by gradient-based minimization algorithm LBFGS. To get out of a local minimum, we perturb all phi/psi angles by a small deviation and then apply LBFGS again to see if a lower energy potential may be reached. We then apply fast relaxation to add side-chain atoms and reduce steric clashes. We generate 20 decoys for each test target and then simply select 5 decoys with the lowest energy as our predictions. We have also tried the Gaussian function for energy potential, but folding with the harmonic function is faster because its derivative is simpler to compute. The harmonic function also improves the average TMscore by about 0.01. For discrete-valued prediction, we convert our predicted discrete distance probability distribution into distance potential using the spline function and then build 3D models by the same protocol.

### Performance metrics

We use precision and F1 value to evaluate contact prediction. To convert discrete probability distributions over the bins to real-valued distance, we compute a weighted average of the distance bins, which is detailed in [5]. To evaluate distance accuracy, we use absolute error, relative error, precision, recall, F1, pairwise distance test (PDT), and high-accuracy pairwise distance test (PHA). For all these metrics, we only consider the distance pairs <15Å. Absolute error is the absolute difference between predicted and native distance while the relative error is the absolute error normalized by the average of predicted and native distance. We measure the recall by the ratio of atom pairs with native distance <15Å that are predicted to have distance <15Å and precision by the ratio of atom pairs with predicted distance <15Å that have native distance <15Å. To calculate PDT and PHA, we first calculate the fraction (R(i)) of predicted distance with an absolute error less than i (i = 0.5, 1, 2, 4, and 8Å). PDT as the average of R(1), R(2), R(4), and R(8) and PHA is the average of R(0.5), R(1), R(2), and R(4).

We use TMScore[18] to evaluate the 3D model quality, which measures the similarity between a 3D model and its experimental structure (i.e., ground truth). It ranges from 0 to 1 and we assume that a 3D model has a correct fold when its TMscore≥0.5.

## Acknowledgments

This work is supported by NIH grant R01GM089753 to J.X.

## Author Contributions

J.L. implemented and tested the code and wrote the manuscript. J.X. conceived the project and revised the manuscript.

## Appendix

**Figure S1.**
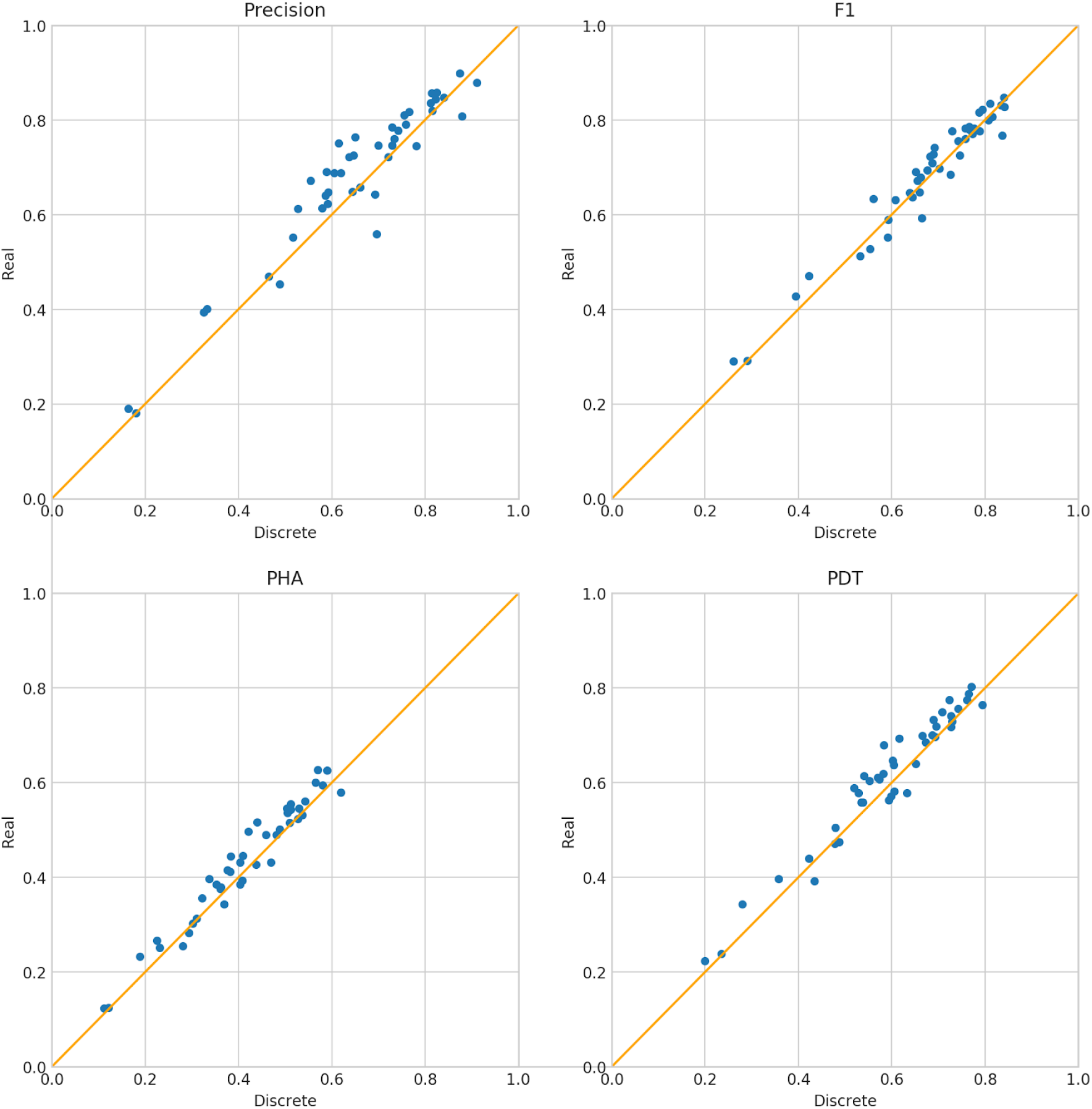
Distance prediction accuracy of our discrete-valued and real-valued method on the 43 FM and FM/BM CASP13 targets. The accuracy is evaluated by precision, F1, PHA and PDT.

**Figure S2.**
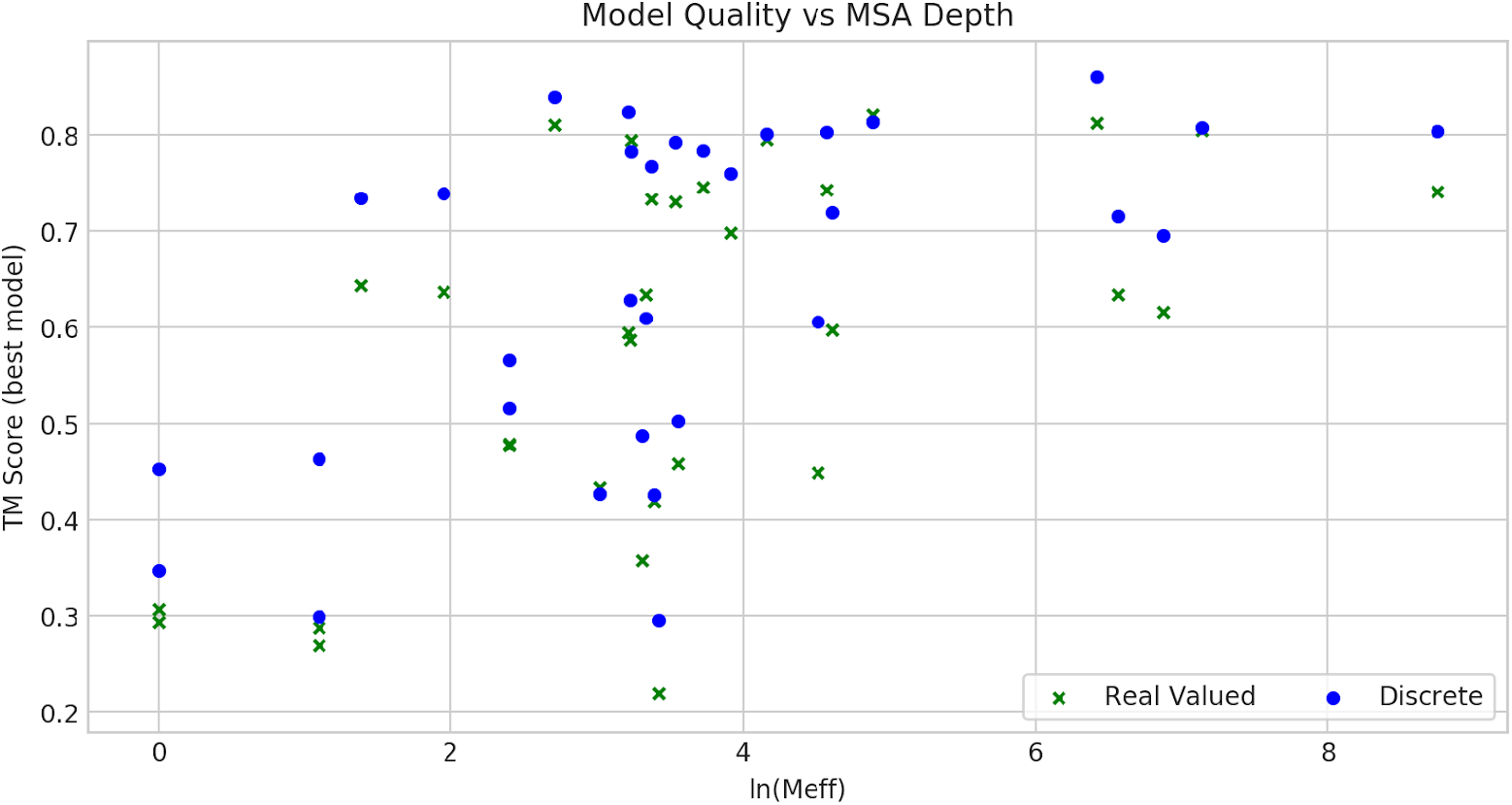
The relationship between the logarithm of MSA depth (i.e., ln(Meff)) and 3D model quality on the 32 CASP13 FM targets.

**Figure S3.**
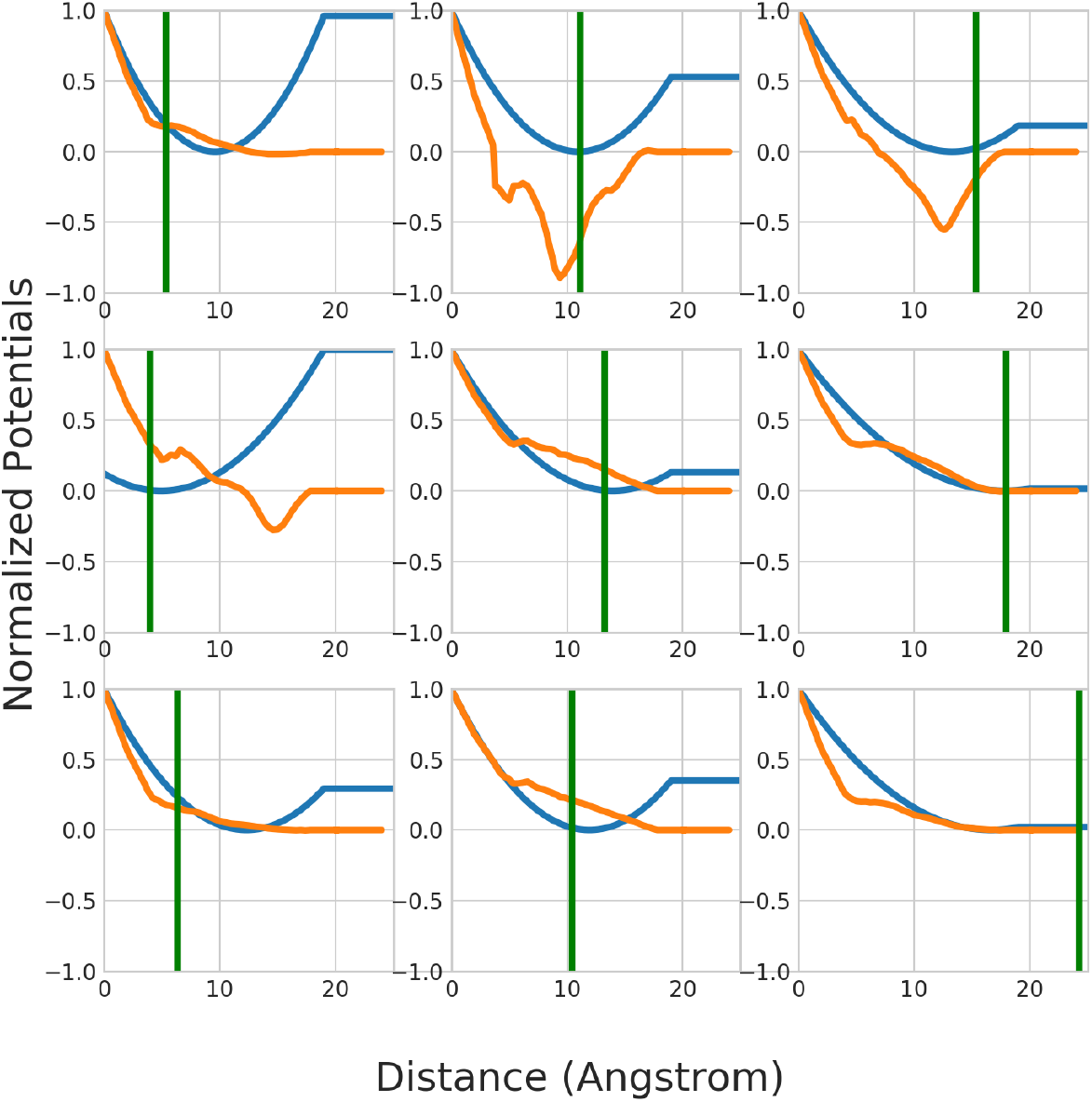
Comparison of predicted distance potentials on one target T1015s1-D1 for short-range (top row), medium-range (middle row), and long-range (last row) residue pairs. Columns correspond to <8Å, 8-15Å, and >15Å. The green line marks the native distance of a residue pair; the blue line corresponds to real-valued prediction and the orange line to discrete-valued prediction.

